# Discovery of potent SARS-CoV-2 nsp3 macrodomain inhibitors uncovers lack of translation to cellular antiviral response

**DOI:** 10.1101/2024.08.19.608619

**Authors:** Alpha A. Lee, Isabelle Amick, Jasmin C. Aschenbrenner, Haim M. Barr, Jared Benjamin, Alexander Brandis, Galit Cohen, Randy Diaz-Tapia, Shirly Duberstein, Jessica Dixon, David Cousins, Michael Fairhead, Daren Fearon, James Frick, James Gayvert, Andre S. Godoy, Ed J. Griffin, Kilian Huber, Lizbé Koekemoer, Noa Lahav, Peter G. Marples, Briana L. McGovern, Tevie Mehlman, Matthew C. Robinson, Usha Singh, Tamas Szommer, Charles W.E. Tomlinson, Thomas Vargo, Frank von Delft, SiYi Wang, Kris White, Eleanor Williams, Max Winokan

## Abstract

A strategy for pandemic preparedness is the development of antivirals against a wide set of viral targets with complementary mechanisms of action. SARS-CoV-2 nsp3-mac1 is a viral macrodomain with ADP-ribosylhydrolase activity, which counteracts host immune response. Targeting the virus’ immunomodulatory functionality offers a differentiated strategy to inhibit SARS-CoV-2 compared to approved therapeutics, which target viral replication directly. Here we report a fragment-based lead generation campaign guided by computational approaches. We discover tool compounds which inhibit nsp3-mac1 activity at low nanomolar concentrations, and with responsive structure-activity relationships, high selectivity, and drug-like properties. Using our inhibitors, we show that inhibition of nsp3-mac1 increases ADP-ribosylation, but surprisingly does not translate to demonstrable antiviral activity in cell culture and iPSC-derived pneumocyte models. Further, no synergistic activity is observed in combination with interferon gamma, a main protease inhibitor, nor a papain-like protease inhibitor. Our results question the extent to which targeting modulation of innate immunity-driven ADP-ribosylation can influence SARS-CoV-2 replication. Moreover, these findings suggest that nsp3-mac1 might not be a suitable target for antiviral therapeutics development.

## Introduction

A strategy for pandemic preparedness is to develop a pipeline of chemical modulators against a wide range of targets in pathogens of pandemic concern, spanning different mechanisms of action, ahead of outbreaks. Coronaviruses are causal agents for multiple outbreaks in recent times, including SARS (2003), MERS (2010), and COVID-19 (2020). Only 3 targets directly implicated in the viral replication cycle have reported chemical inhibitors that translate to antiviral activity: Numerous approved therapeutics and clinical candidates to date target the main protease and RNA-dependent RNA polymerase (*1*), while recent work reported potent antiviral leads against the papain-like protease (*2, 3*). Beyond proteins that are directly responsible for viral replication, an emerging body of literature elucidates the role of viral proteins which counteract the host immune response in causing virulence and pathogenesis. Finding chemical inhibitors against these targets, thus modulating the competition between infection and immunity, is an underexplored mechanism of action for antivirals.

SARS-CoV-2 nsp3-mac1 is a macrodomain with ADP-ribosylhydrolase activity (see **Figure 1A**) which is hypothesised to modulate host immune response (*4*). Upon infection, interferon-mediated immune response is initiated by host cells, leading to the expression of poly-(ADP-ribose)-polymerases (PARPs). PARPs catalyse the post-translational addition of ADP-ribose to a large range of proteins triggering degradation, which nsp3-mac1 enzymatically reverses (*5–7*). Macrodomains are conserved across coronaviruses, and also found in togaviridae and hepeviridae (*6*), suggesting that it may be an integral part of the evolutionary tug-of-war between infection and immunity.

**Figure 1:**
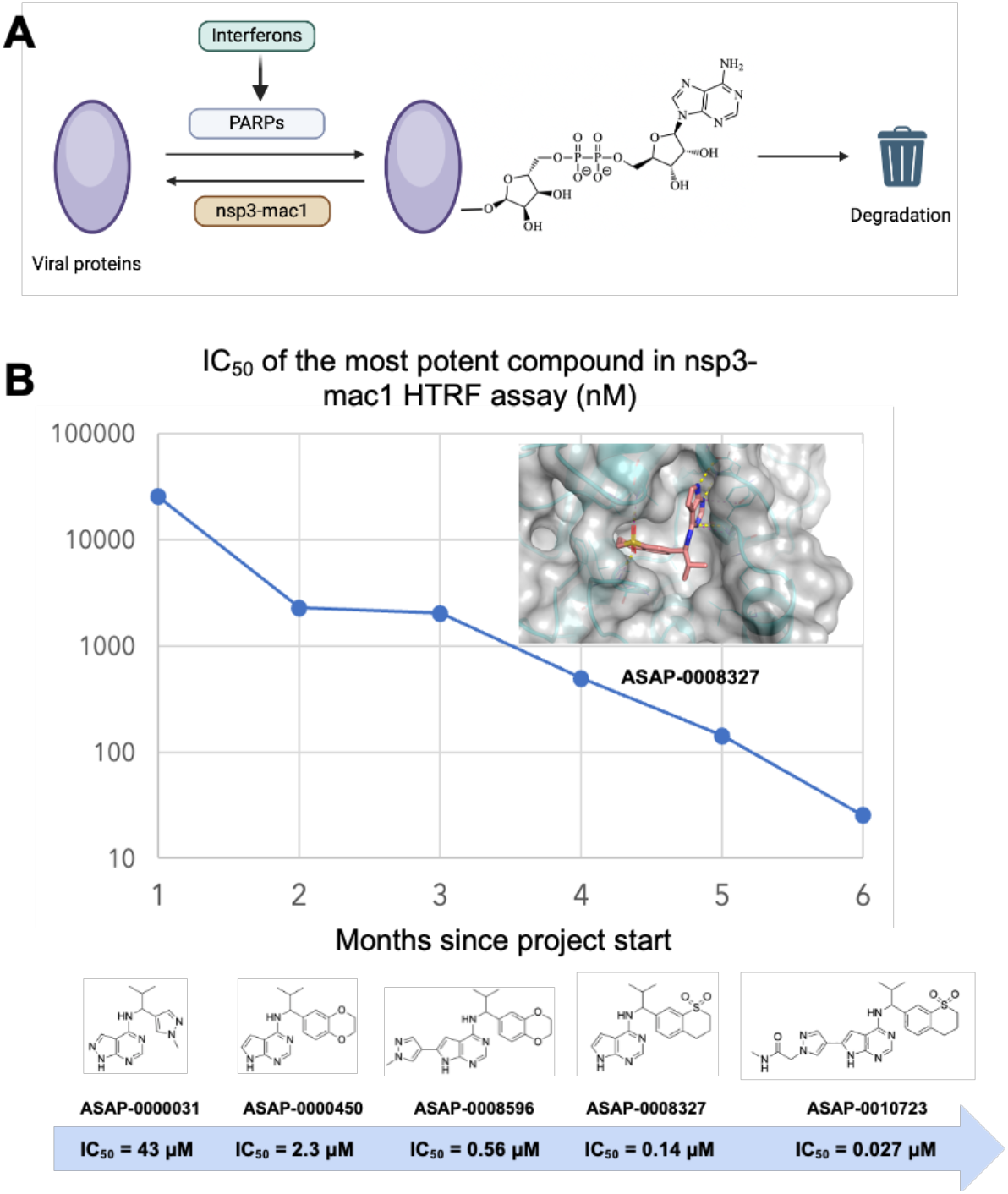
Summary of the SARS-CoV-2 nsp3-mac1 campaign. (A) Schematic of the role of viral nsp3-mac1 in reversing ADP-ribosylation via innate immune response. (B) We report a rapid fragment-to-lead campaign, realising significant strides in potency. The inset shows the structure of a lead compound (ASAP-0008327), confirming active site engagement (PDB ID 7GZU).

Recent reverse genetics studies on SARS-CoV-2 show that abrogation of nsp3-mac1 enzymatic activity causes a decrease in viral replication in cell models, though the effect depends on cellular immune response and the effect is notably absent in VeroE6 (*8, 9*). In a mouse *in vivo* efficacy model, nsp3-mac1 mutants abrogate pathogenesis of SARS-CoV-2 (*8, 9*), inline with previous studies on SARS-CoV (*6*). However, recent studies on murine hepatitis virus (MHV) demonstrate a difference in phenotype between macrodomain’s enzymatic function and mono-ADP-ribose:nsp3-mac1 binding (*10*), corroborating the literature on alphavirus macrodomain (*11, 12*). A chemical biology approach of probing the phenotypic effect of orthosteric competitive inhibitors, which abrogate enzymatic function as well as occlude the binding site in a concentration-dependent manner, could reveal complementary insights to reverse genetics studies.

During the COVID-19 pandemic, a crystallographic fragment screening campaign was published against nsp3-mac1, showing a dense assortment of over 234 fragment hits which densely decorate the active site (*13*). Building on the fragment hits, which have weak biochemical activity, a structure-guided iterative medicinal chemistry campaign furnished inhibitors with measurable nsp3-mac1 activity by merging fragments (*14*). However, no antiviral activity was reported for that chemical series, and significant opportunities abound for improving the cell permeability and potency. Parallel effort starting from high throughput screen furnished inhibitors with modest activity against nsp3-mac1 and reported measurable antiviral activity in combination with interferon gamma (*15*).

Herein we report a fragment-based lead discovery campaign (see **Figure 1B** for a summary) which furnished cell-permeable leads with low nM nsp3-mac1 activity. Using our tool compounds, we show that whilst nsp3-mac1 inhibition leads to an increase in ADP ribosylation, it does not translate to cellular antiviral activity in iPSC-derived pneumocytes nor cell culture systems. Further, there’s no evidence of synergy between nsp3-mac1 inhibition and Mpro or PLpro inhibition, nor interferon gamma treatment. Taken together, our work questions the extent to which targeting nsp3-mac1’s role in innate immunity-driven ADP-ribosylation modulates SARS-CoV-2 replication.

### Pharmacophore inference generates hits from fragments

The starting point of this lead discovery campaign was a high-throughput crystallographic fragment screen, furnishing 234 fragment hits after screening 2593 diverse fragments. These hits densely cover the enzyme active site where ADP-ribose binds (**Figure 2A**). However, the challenge with fragment starting points is that they have weak to no affinity (*14*). To overcome this, we exploit the density of fragment hits via a statistical approach rather than focusing on individual fragments.

**Figure 2:**
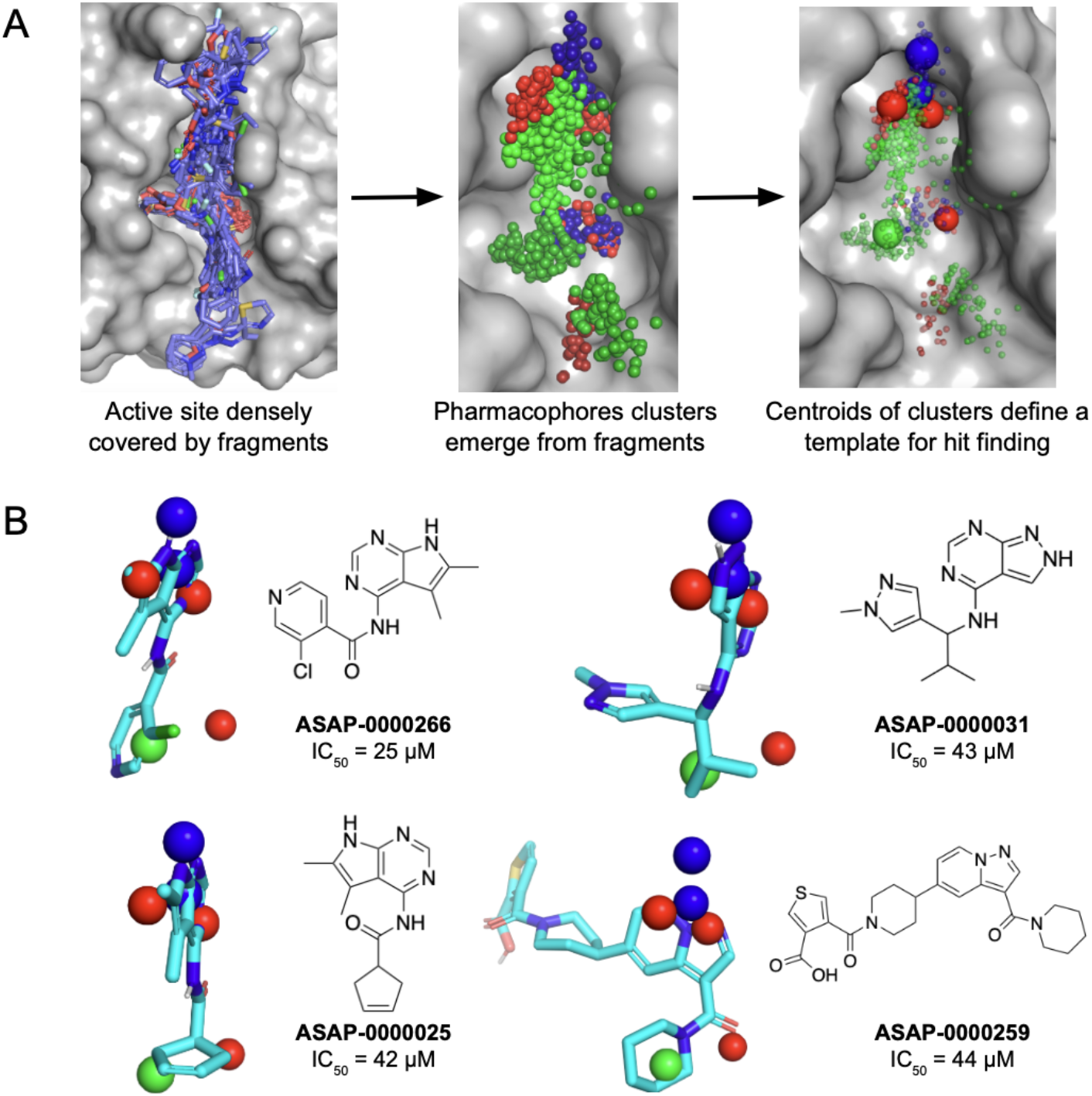
Pharmacophore identification enables fragment-based hit discovery. (A) Conversion of fragments into pharmacophore clusters. The red, blue and green spheres correspond to hydrogen bond acceptor, donor and hydrophobic pharmacophores, respectively. (B) Selected hit molecules superimposed on the identified pharmacophores.

We coarse-grain fragments into pharmacophores -- chemical motifs which could lead to specific protein-fragment interactions -- and use the statistics of pharmacophores to infer salient interactions that would lead to potency. Our hypothesis is that although individual fragments are weak binders, if multiple fragments hit the same location, with the same pharmacophore, that pharmacophore is likely to be salient to binding. Further, we can correlate the strength of that interaction by the number of times that a pharmacophore is found, across different fragments, in a particular location of the binding site. Once we identify salient pharmacophores, we can search purchasable molecules which fit the pharmacophores and share chemotypes of the fragment hits. By focusing on purchasable molecules with pharmacophores which match the fragments hits, we hope to avoid the challenging chemistry that is often associated with chemically introducing handles for growing individual fragments (*16*), whilst mitigating strain penalty that is often incurred from directly linking fragments (*17*). Our approach combines pharmacophore-based lead identification (*18–20*) with statistical approaches of coarse-graining fragment hits into pharmacophore fields (*21*).

**Figure 2A** depicts the fragment hits as a pharmacophore field. For ease of visualisation, each pharmacophore is represented as a Gaussian distribution, and we perform clustering to locate local maxima which correspond to spatial regions which are enriched with a particular pharmacophore. We search for compounds in Enamine REAL, a virtual library of 36 billion made-on-demand molecules, and rank by goodness of match to the pharmacophores, and we experimentally screened a diverse subset of the matches. **Figure 2B** shows selected hits with associated nsp3-mac1 activity.

### Lead discovery with synthesis-guided design

We identified **ASAP-0000031** as a ligand-efficient starting point for further optimisation using an iterative high throughput library chemistry approach and computational building block selection.

We computationally overlaid **ZINC000089254160_N3 (PDB ID 5RSJ)** with **ASAP-0000031** (**Figure 3A**), which revealed a vector for elaboration. Based on that, we enumerated a library based on nucleophilic aromatic substitutions (SNAr), with the core replaced to pyrrolopyrimidine due to reagent availability. We specifically focused on secondary amines with heteroaromatic groups connected at the alpha position. In total, 42 compounds were made. A histogram of pIC_50_ reveals that although most compounds were inactive, some were breakthrough outliers in potency, most notably **ASAP-0000450 (Figure 3)**.

**Figure 3:**
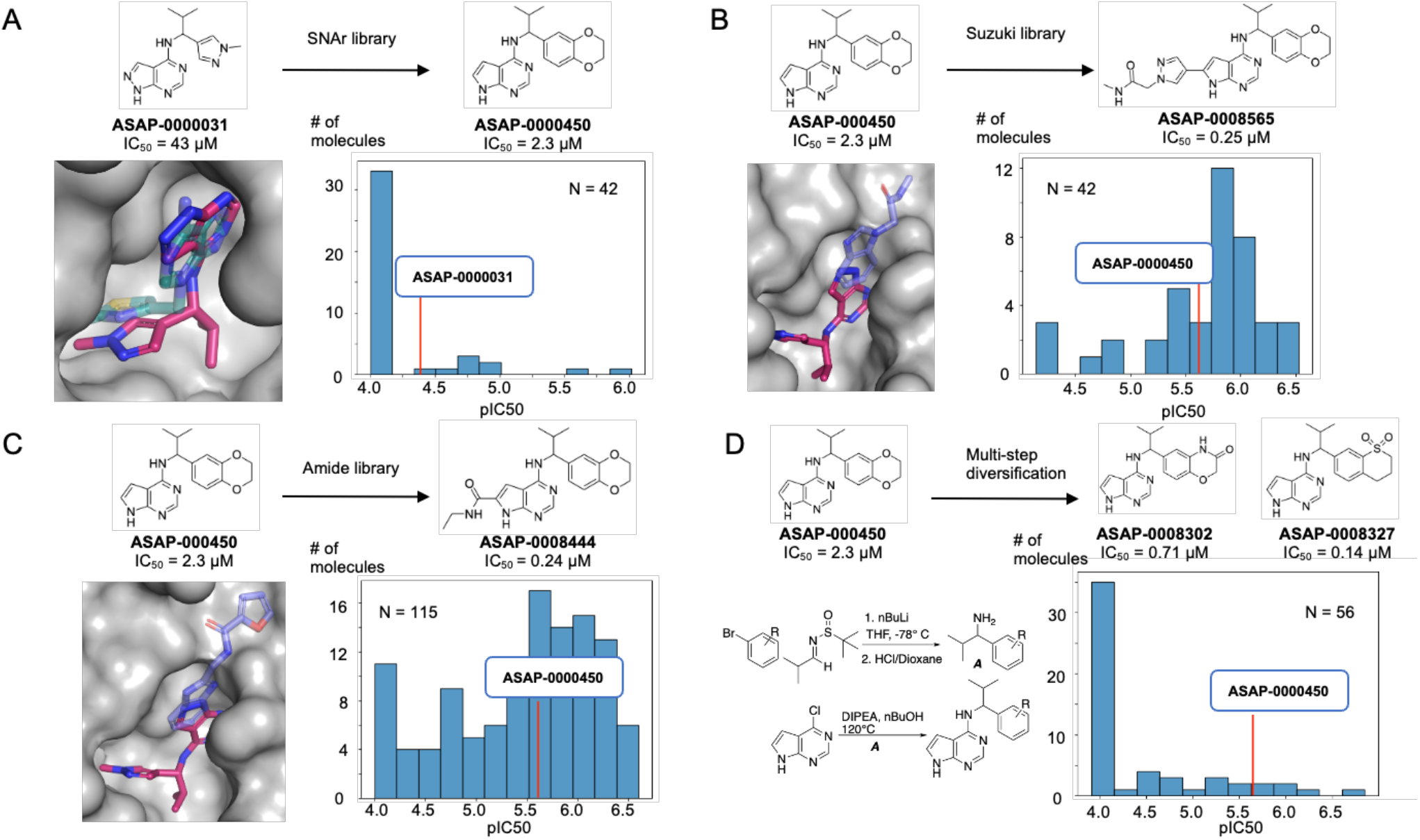
Hit to lead chemistry via library chemistry. We designed parallel medicinal chemistry libraries based on (A) SNAr, (B) Suzuki, and (C) Amide chemistries to realise step change in potency. (D) shows a multistep library which enables variations in the benzodioxane motif. Computational overlays between crystallographic structure of fragments (transparent) and our lead compounds (solid) provided inspiration for library design.

To follow up on **ASAP-000450**, we first elaborated the pyrrolopyrimidine core. The computational overlay between **ASAP-0000031**, the fragments **EN300-697611 (PDB ID 5S24)** (**Figure 3B**), and **Z26781964 (PDB ID 5S2T)** (**Figure 3C**) suggested a library based on Suzuki coupling from the 6-bromo pyrrolopyrimidine, and amide coupling library using a building block with a carboxylic acid at the 6 position of pyrrolopyrimidine. These libraries were significantly more promising, with most compounds leading to an improvement in potency. The champion compounds were **ASAP-0008565** and **ASAP-0008444**, both 10x increases in potency relative to the starting point.

We next turn to explore additional vectors, aiming to obtain synergistic gains with the aforementioned exploration of the pyrrolopyrimidine core. A promising region for further optimization is the benzodioxane motif, as the SNAr library (**Figure 3A**) was limited in suitable amine building blocks. Hence we broadened our library approach to consider multi-step syntheses, leveraging the broad availability of aryl bromide building blocks to install benzodioxane modifications in the first step. This set of compounds was considerably more challenging, both synthetically (scheme shown in **Figure 3D**) and also potency-wise. Most compounds were inactive, but demonstrable improvements abound with **ASAP-0008302** and **ASAP-0008327**, showing IC50s of 0.71 and 0.14 μM, respectively (Figure 3D) .

We then focused on combining potency improvements from each vector, whilst using predicted *in vitro* physicochemical and ADME properties to prioritise design from the vast combinatorial space. To deliver a tool compound, we specifically focus on high solubility and permeability, whilst maintaining moderate human and rodent liver microsome clearance. **Figure 4A** shows selected structure-activity relationships. Variations in isopropyl moiety R_1_ show whilst polar groups are not tolerated, increasing lipophilicity does not lead to significant potency improvements. From the benzodioxane end R_2_, a rigid cyclic scaffold is required for potency, whilst significant potency can be unlocked by increasing polarity and hydrogen bond modulation. The structure-activity relationship is synergistic (**Figure 4B**), and **Figure 4C** shows that potency gains are achievable whilst maintaining solubility and permeability.

**Figure 4:**
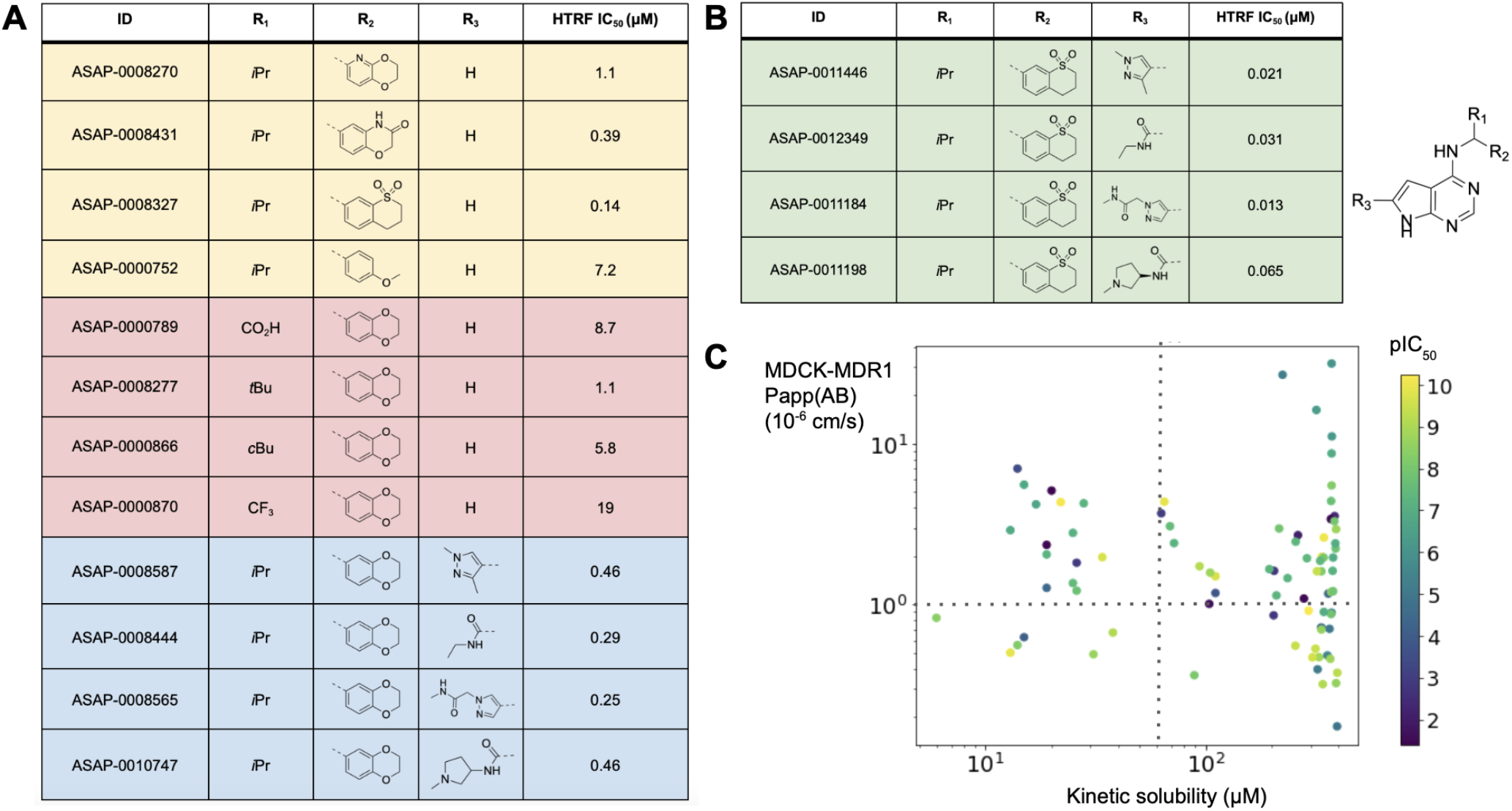
Combining structure-activity relationship furnishes potent inhibitors. (A) shows key variations in one substituents around the scaffold (potency values are reported for the racemic mixture). (B) shows that the structure-activity relationship is synergistic; we report the IC_50_ of the bioactive pure enantiomer. (C) shows potency pIC_50_ as a function of permeability and solubility, revealing no inherent tradeoff. The dotted lines demarcate regions of high permeability (>10^-6^ cm/s) and solubility (>50 μM).

### Structural biology reveals salient interactions of lead compounds

Our chemistry efforts were supported by high-throughput crystallography, screening a total of 890 compounds which resulted in the release of 90 ligand-bound structures (PDB group deposition ID: G_1002283). Initially, a P4_3_ crystal form was used for compound soaking, based on the previous crystallographic fragment screen of SARS-CoV-2 nsp3-mac1, as this crystal form offered high resolution data and good accessibility to the ADPr binding site (*13*). However, scaffold expansion of the pyrrolopyrimidine core required the establishment of a second crystal system (P1 2_1_ 1) to enable co-crystallization of the lead compounds with nsp3-mac1 due to the occlusion of ligand binding by crystal contacts present in the P4_3_ system.

The top lead compounds **ASAP-0011198, ASAP-0011446**, and **ASAP-0011184** share a common core structure and, therefore, highly similar binding poses (**Figure 5A-C**). The sulfone containing scaffold (R_2_) bound to the phosphate subsite of nsp3-mac1 forming hydrogen bond interactions with the backbone nitrogens of Val49 and Ile131, and via a water bridge with Ala38, as well as hydrophobic interactions with the side chains of Phe132, Ile131, Val49 and Ala38. The isopropyl group (R_1_) forms hydrophobic interactions with the side chains of Asp157 and Leu160. The pyrrolopyrimidine core sits in the adenine site of nsp3-mac1, tightly bound by multiple hydrogen bonds with Asp22, Ile23 and Phe156. Additionally, the Phe156 residue forms key interactions with the pyrrolopyrimidine via hydrophobic and π-stacking interactions. The scaffold extension (R_3_) of the pyrrolopyrimidine core exhibits few specific interactions with nsp3-mac1, such as hydrophobic interactions with Ala52 or, in the case of the amide moiety in **ASAP-0011198**, hydrogen bond interactions with Asp22.

**Figure 5:**
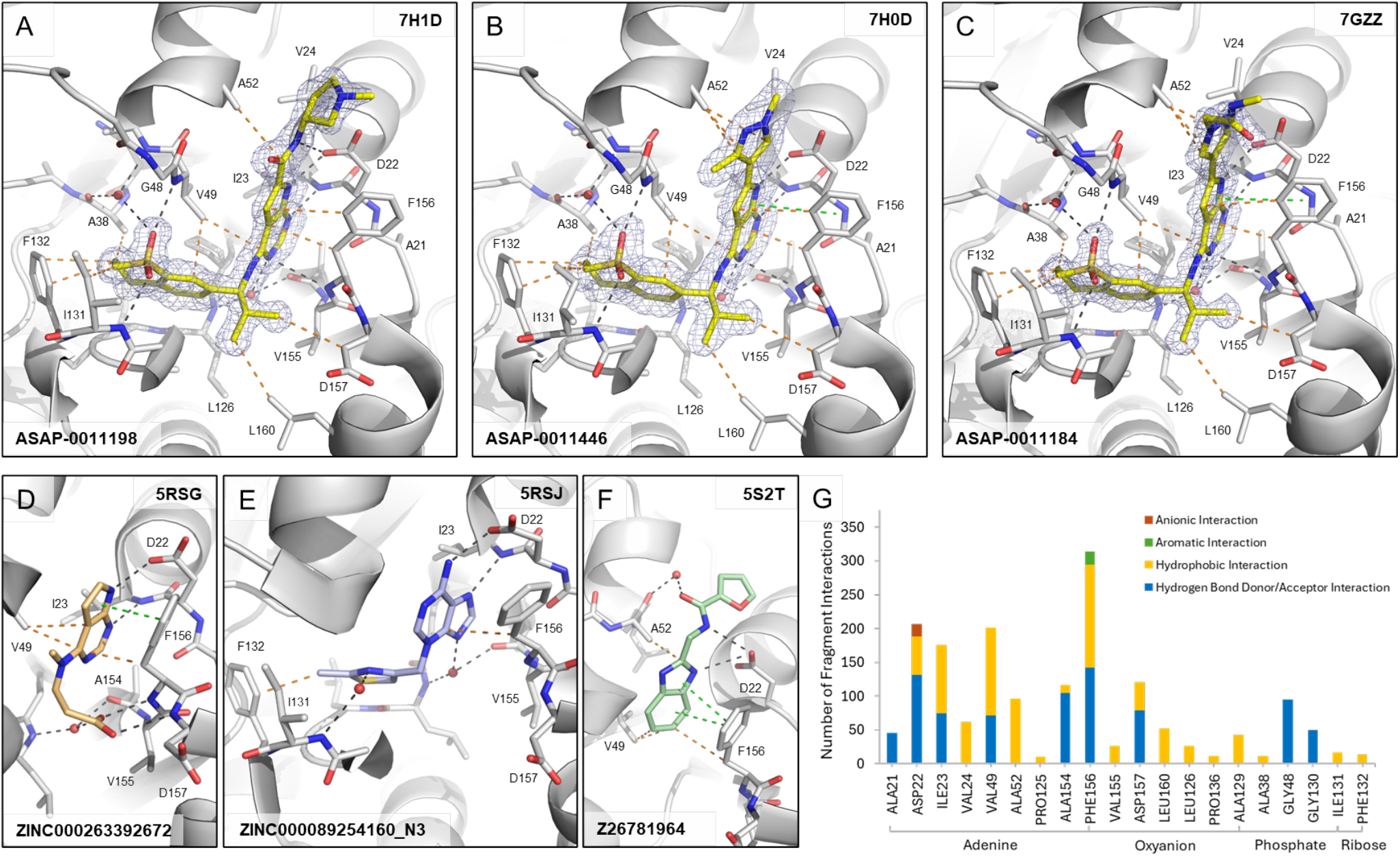
Binding interactions of lead compounds and fragments against SARS-CoV-2 nsp3-mac1. Crystal structures of (A) **ASAP-0011198 (PDB ID 7H1D)**, (B) **ASAP-0011446 (PDB ID 7H0D)** and (C) **ASAP-0011184 (PDB ID 7GZZ)** bound to the ADPr-binding site of nsp3-mac1 and selected fragments (D) **ZINC000263392672 (PDB ID 5RSG)**, (E) **ZINC000089254160_N3 (PDB ID 5RSJ)** and (F) **Z26781964 (PDB 5ST2)** that were used as inspiration for the lead compounds. Dashes depict polar (black), aromatic (green), and hydrophobic (orange) interactions between ligands and nsp3-mac1. 2Fo-Fc map (σ=1.0) is displayed as blue mesh around the lead compounds. (G) Analysis of interactions by all fragment hits found in the crystallographic fragment screen against nsp3-mac1 (13).

The salient interactions formed by our tool compounds were observed in the original crystallographic fragment screen of SARS-CoV-2 nsp3-mac1 (*13*). Enumeration of all interactions formed by the fragment hits (**Figure 5G**), illustrates that our lead compounds recapitulate many of the common interactions observed in each respective subsite, with the exception of hydrogen bond interactions with backbone nitrogens of residues Phe156 and Asp157 in the oxyanion hole. Unsurprisingly, these salient interactions were observed in the fragments (**ZINC000263392672, ZINC000089254160_N3** and **Z26781964**) which inspired our library design and elaborations (**Figure 5D-F**).

### Tool compounds reveal a lack of translation from mac1 inhibition to antiviral activity

Figure 6. shows that our top leads have drug-like solubility, permeability and lipophilicity, which are crucial figures of merit for a cellular tool compound. In terms of metabolic stability, **ASAP-0011198** has low clearance, whereas **ASAP-0011446** and **ASAP-0011184** have moderate-to-high clearance; clearance is not a key property we focused on in the development of *in vitro* tool compounds, and taken together the data suggests that there is no scaffold-wide liability in metabolic stability. We derisk potential for off-target activity by profiling the compound in a CEREP panel, a standard secondary pharmacology assessment, as well as a kinase panel to derisk a potential hinge-binding motif in our leads, and a PARP panel to derisk activity against human ADP polymerase. Reassuringly, the compounds are selective, with the only off-targets being moderate human CHRM2 and KOR receptors activity associated with **ASAP-0011198**.

Profiling the 3 tool compounds against SARS-CoV-2 infection in HeLa-ACE2 cells reveals no detectable antiviral effects.This leads to the salient conclusion of this work, that nsp3-mac1 inhibition did not translate to antiviral activity for this series in a cell line amenable to a high-throughput screening campaign. We next consider whether nsp3-mac1 inhibition can lead to antiviral activity in combination with established SARS-CoV-2 antivirals. As such, we further profile the compounds in combination with the measured EC50 concentration of nirmatrelvir, a MPro inhibitor, and GRL0617, a PLpro inhibitor. No potentiation of nsp3-mac1 antiviral activity is detected in combination with these well established protease inhibitors.To attribute the lack of antiviral activity to the insufficiency of ADP-ribosylation modulation through mac1 inhibition, we confirm target engagement via cellular thermal shift assay (**Figure 7A**), and increase of ADP-ribosylation in cells upon exposure to our compounds. **Figure 7B** shows that **ASAP-0011184** increases ADP-ribosylation in infected HelaACE-2 cells, in line with the biochemical hypothesis.

**Figure 6:**
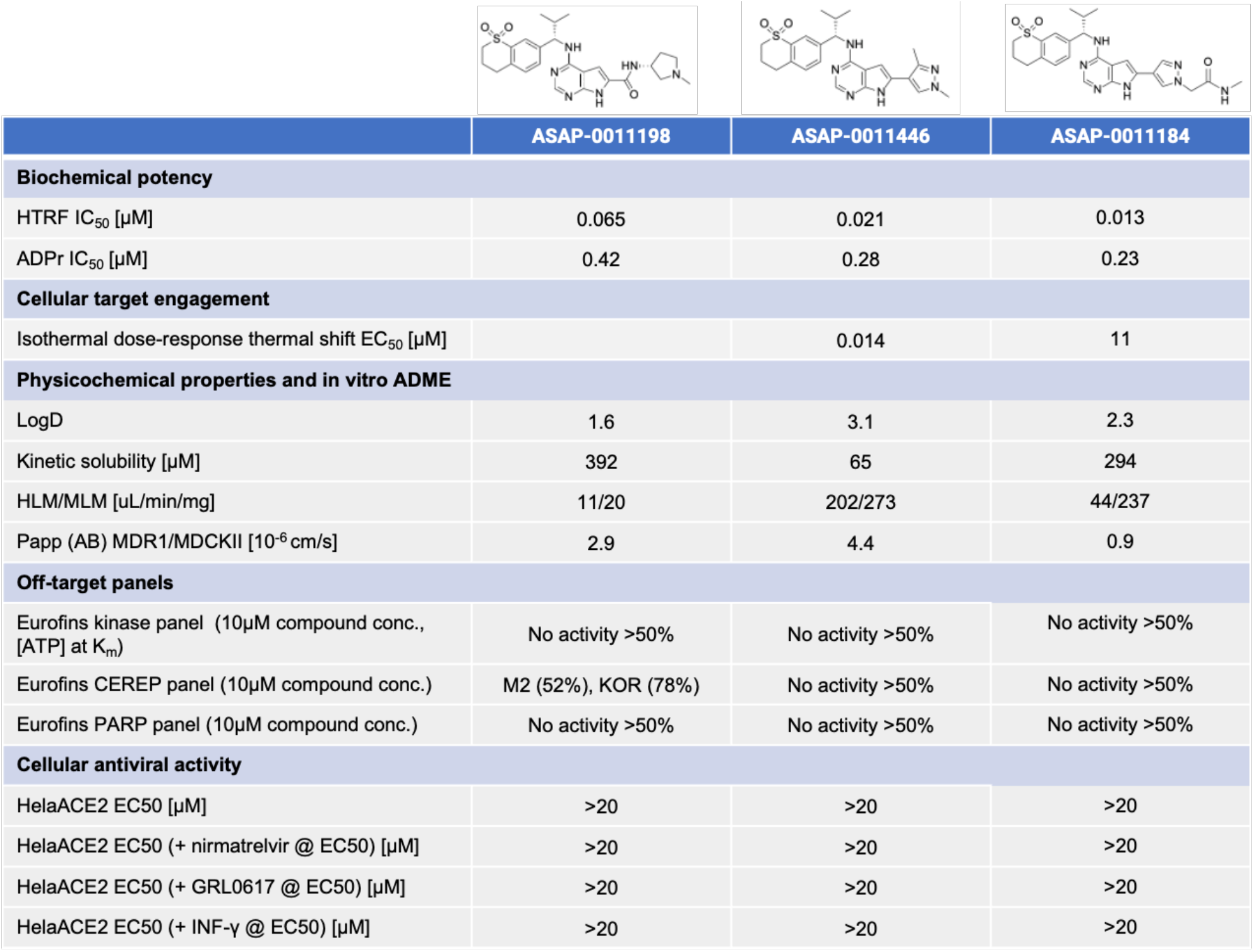
Our campaign furnished a toolbox of chemical probes against SARS-CoV-2 nsp3-mac1. The table shows key physicochemical, in vitro ADME, and selectivity data for tool compounds. These tool compounds clearly reveal that nsp3-mac1 inhibition does not translate to antiviral activity.

**Figure 7:**
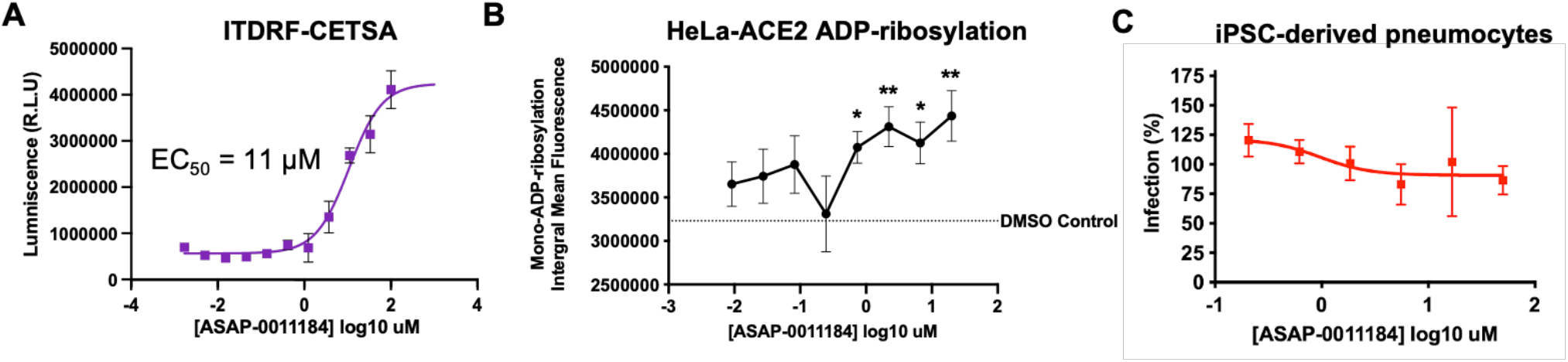
nsp3-mac1 inhibition does not translate to antiviral response. (A) Cellular Thermal Shift Assay (CETSA) demonstrating cellular target engagement. (B) Mono-ADP-ribosylation integral mean fluorescence of infected HelaACE2 cells as a function of compound concentration, showing a statistically significant increase upon dosing **ASAP-0011184**. (C) No statistically significant antiviral effect is observed for **ASAP-0011184** in an iPSC-derived pneumocyte system. Data was analysed by two-way ANOVA (^*^P < 0.05, ^**^P < 0.01, ^***^P < 0.001, and ^****^P < 0.0001).

Finally, we consider the hypothesis that interferon response triggers ADP-ribosylation, thus profiling the compounds in enhanced and physiologically relevant interferon environments could reveal detectable antiviral effects. Compounds were tested in combination with measured antiviral EC_50_ concentration of IFN-γ, and nsp3-mac1 inhibition still does not translate to antiviral activity even in this enhanced interferon response setting (**Figure 6**). Furthermore, HeLa-ACE2 cells, and all cancerous cell lines, display defective innate immune response (*1*). As such, we also assess our compound in a primary-like iPSC-derived pneumocyte system (**Figure 7C**), showing likewise a lack of antiviral activity in this physiologically relevant innate immunity system.

## Discussion

Herein we report a fragment-to-lead campaign directed at SARS-CoV-2 nsp3-mac1, arriving at potent and selective tool compounds with drug-like physicochemical and ADME properties. We show that inhibition of macrodomain activity increases ADP-ribsolyation, but surprisingly does not abrogate viral replication in cellular systems.

Our study is limited in two important aspects. First, we cannot rule out whether further increase in potency of our inhibitors will eventually lead to cellular antiviral activity. From mutational studies, complete suppression of macrodomain function does impact viral replication in cellular systems (*8, 9*). However, it is questionable whether an inhibitor can phenocopy the complete suppression of enzyme function. Approaches such as targeted protein degradation might be more promising, and we hope our inhibitors, with structural information, could provide a starting point for the design of degraders. We note that our results are in tension with ref (*15*), which reported measurable cell potency even with nsp3-mac1 inhibitors with modest potency; our inhibitors are significantly more potent and were subjected to broad target selectivity panels to ensure off-target effects are not confounding.

Second, our study is limited to cellular antiviral activity. The literature suggests that the effect of macrodomain inhibition might be more pronounced in *in vivo* models, with a rich repertoire of immune responses, than in cellular models (*8, 9*). However, the clinical translatability of the immune response in animal models of infection is still an open question. In fact, the development path of current SARS-CoV-2 antivirals relied on human primary cell models as the estimate of potency, whilst animal efficacy has been viewed as corroboratory evidence (*1*). We hope our inhibitors will contribute to future studies which might reveal the translatability of immune response in animal models of SARS-CoV-2 infection.

In line with this goal, the ASAP Discovery Consortium is releasing extensive biochemical, antiviral, and ADMET data from this drug discovery program on the over 1000 compounds tested as part of the nsp3-mac1 inhibitor discovery efforts (appended in the Supplementary Materials). Furthermore, as part of our open science mission, we have already deposited extensive assay data from this program into robust molecular databases such as ChEMBL (*22*) for streamlined, upcoming release to the drug discovery community. Recognizing that these results call into question the feasibility of pursuing an actively worked on target, we hope this comprehensive data package proves helpful for further investigation.

## Methods

**HTRF Mac1 assay** https://dx.doi.org/10.17504/protocols.io.eq2ly7r2mlx9/v3

### ADPr hydrolase assay

ADPr hydrolase activity assay was conducted in 96-well plate with pre-dispensed compounds in 8-point dose response starting from 4 μM (control wells were backfilled with DMSO). Assay was conducted in 96-well plate with pre-dispensed compounds in 8-point dose response starting from 4 μM (control wells were backfilled with DMSO). Nsp3-mac1 200 nM in final assay concentration was added to all wells besides “NAD+ alone” well. The plate was centrifuged at 1,500 rpm for 1 min, and then Incubated at room temperature for 15 min. After incubation, αNAD+ in final concentration of 8 μM were added to each well, and the plate was centrifuged again for 1 min. Then, the plate was incubated under shaking at room temperature for 2.45 h, followed by freezing at -80°C until LC-MS/MS analysis.

A 30-μl sample aliquot was mixed with 90 μl of methanol and 1μl of 50 μM 15N5-AMP (an internal standard). The mix was shaken for 10 min, then centrifuged (∼21,000 g) for 5 min to precipitate the protein. The supernatant was placed into a 250-uL insert of LC-MS vial for the following analysis.

The LC-MS/MS instrument consisted of Acquity I-class UPLC system and Xevo TQ-S triple quadrupole mass spectrometer (Waters). Chromatographic separation was done on a 100 x 2.1-mm i.d., 1.7-μm Atlantis Premier BEH C18 AX column (Waters) with mobile phases A (20mM ammonium formate, pH 3.0) and B (acetonitrile) at a flow rate of 0.3 ml/min and column temperature 25°C. A gradient was as follows: 2-min interval the column was held at 2% B, then during 5-min linear increase till 100% B, followed by 1-min held at 100% B, then 1-min back to 2% B and equilibration at 2% B for 1 minute. Samples kept at 8°C were automatically injected in a volume of 1 μl.

Mass detection was carried out using electrospray ionization in the positive mode. Argon was used as the collision gas with flow 0.10 ml/min. The capillary voltage was set to 2.5 kV, source temperature - 120°C, desolvation temperature - 600°C, desolvation gas flow - 900 L/min, cone voltage 16V. Analytes were detected using multiple reaction monitoring (MRM, m/z) with the following precursor and fragment masses at selected collision energy (eV, indicated in brackets): 560 > 136 (34) and 560 > 348 (16) for ADPR, 664 > 524 (15) and 664 > 541.7 (15) for αNAD+, 353.1 > 97 (28) and 353.1 > 141 (20) for 13N5-AMP (IS) m/z. Standard curve of the ADPR and αNAD+ mix in the range of 0.01-10 uM was used. MassLynx and TargetLynx software (v.4.2, Waters) were used to acquire and analyze the data.

### Cells and Viruses

HeLa-ACE2 cells (BPS Bioscience, were maintained in DMEM (Corning) supplemented with 10% FBS, 0.5 μg/mL puromycin, and penicillin/streptomycin (Corning) at 37°C and 5% CO2. All cell lines used in this study were regularly screened for mycoplasma contamination using the MycoStrip™ - Mycoplasma Detection Kit (InvivoGen, rep-mys-20). Cells were infected with SARS-CoV-2, isolate USA-WA1/2020 (BEI Resources NR-52281). Viruses were grown in Vero-TMPRSS2 cells (BPS Bioscience) for 4–6 d; the supernatant was clarified by centrifugation at 4,000g for 5 min and aliquots were frozen at −80°C for long term use. Expanded viral stocks were sequence-verified to be the identified SARS-CoV-2 variant and titered on Vero-TMPRSS2 cells before use in antiviral assays. Infections with viruses were performed under biosafety level 3 (BSL3) containment in accordance to the biosafety protocols developed by the Icahn School of Medicine at Mount Sinai.

### Antiviral assay

Four thousand HeLa-ACE2 cells (BPS Bioscience) were seeded into 96-well plates in DMEM (10% FBS) and incubated for 24 hours at 37°C, 5% CO2. Two hours before infection, the medium was replaced with 100 μL of DMEM (2% FBS) containing the compound of interest at concentrations 50% greater than those indicated, including a DMSO control. Plates were then transferred into the BSL3 facility and 100 PFU (MOI = 0.025) was added in 50 μL of DMEM (2% FBS), bringing the final compound concentration to those indicated. Plates were then incubated for 24 hours at 37°C. After infection, supernatants were removed and cells were fixed with 4% formaldehyde for 24 hours prior to being removed from the BSL3 facility. The cells were then immunostained for the viral N protein (an inhouse mAb 1C7, provided by Dr. James Duty) with a DAPI counterstain. Infected cells (488 nm) and total cells (DAPI) were quantified using the Cytation 1 (Biotek) imaging cytometer. Infectivity was measured by the accumulation of viral N protein (fluorescence accumulation). Percent infection was quantified as ((Infected cells/Total cells) - Background) *100 and the DMSO control was then set to 100% infection for analysis. Data was fit using nonlinear regression and IC_50_s for each experiment were determined using GraphPad Prism version 10.0.0 (San Diego, CA). Cytotoxicity was also performed using the MTT assay (Roche), according to the manufacturer’s instructions. Cytotoxicity was performed in uninfected cells with same compound dilutions and concurrent with viral replication assay. All assays were performed in biologically independent triplicates.

Combination experiments were performed by first determining the EC50 of nirmatrelvir, GRL-0617, or IFN-γ alone, as described in the paragraph above. Then, the determined EC50 concentration of each drug was added to all control and experimental wells of a plate and a dose-response of the Mac1 inhibitors was added to the on top of each of these 3 treatments. All assays were performed in biologically independent triplicates.

### Tracking ADP-ribosylation

8,000 HeLa-Ace-2 cells were seeded into 96-well plates in 100 μl of growth media (DMEM, 10% FBS, 1% 100x Penicillin & Streptomycin, .2% Plasmocin, .006% Puromycin and 1% 100x Nonessential Amino Acids) and were incubated for 24 hours at 37°C and 5% CO2. Cells were seeded with a viability ≥ 80% as determined through Trypan Blue staining and analysis by the Countess 3 Automated Cell Counter (ver. 1.2.881.580). After 24 hours the growth media was aspirated from the well plates and replaced with 100 μl of infection media (DMEM, 2% FBS, 1% 100x Penicillin & Streptomycin, 1% Nonessential Amino Acids, and 1% HEPES). The drug treatment was administered to treatment wells through direct dilution by the Tecan D300e (ver. 3.4.3) at a concentration of 30 μM (50% higher than the final concentration). DMSO concentrations between drug dilutions and controls were additionally normalized to the highest concentration of DMSO through direct dilution by the Tecan D300e. Following drug treatment, the well plates were incubated for 2 hours at 37°C and 5% CO2 before infection. Treatment and DMSO control wells were infected with 50 μl of WT SARS-CoV-2/WA1 at an MOI of 0.1, with uninfected wells receiving a mock infection of 50 μl infection media, diluting the drug to the final concentration of 20 μM. Subsequently after infection, the cells were incubated at 37°C and 5% CO2 for 24 hours, before being fixed at a final solution of 4% formaldehyde overnight at 4°C.

ADP-Ribosylation Immunostaining was performed through incubating fixed cells in a 50 μl solution of primary Ab targeting SARS-CoV-2 NP protein (an inhouse mAb 1C7C7, provided by Dr. Andrew Duty) and Mono-ADP ribosylation (rAb MABE1076) at a ratio 1:1000 in 1% BSA-PBS, overnight at 4°C. After two washes with 0.1% Tween20-PBS, the cells were incubated in a 50 μl solution of secondary Ab containing anti-mouse Alexa fluor 488 and anti-rabbit Alexa fluor 647 at a ratio of 1:1000 in 1% BSA-PBS, with a DAPI counterstain for 1 hour at RT. Prior to fluorescent imaging, cells were washed twice with 0.1% Tween20-PBS before being imaged in 1x PBS. Fluorescent images were captured and quantified using BioTek Cytation10. Mono ADP ribosylation was measured by the integral mean fluorescence of 594/633 nm cells. Infected and uninfected cell populations were distinguished through secondary gating using 488 nm mean fluorescence. ADP-ribosylation measurements for drug treatments were captured at a n = 9; DMSO, n = 16; uninfected n = 8. Cellular analysis was performed using Gen5.

### Isothermal dose response thermal shift assay

A PCR product encoding SARS Cov-2 macrodomain residues 2 to 176 was cloned into the XhoI and BamHI restriction sites of the pBIT3.1-N vector (Promega) which has an N-terminal 11 amino acid HiBiT tag.

PBIT NHIBIT SARS-CoV-2 Macrodomain

M**VSGWRLFKKIS***GSSGGSS*<u>PEVNSFSGYLKLTDNVYIKNADIVEEAKKVKPTVVVNAANVY</u> <u>LKHGGGVAGALNKATNNAMQVESDDYIATNGPLKVGGSCVLSGHNLAKHCLHVVGPNVNK</u> <u>GEDIQLLKSAYENFNQHEVLLAPLLSAGIFGADPIHSLRVCVDTVRTNVYLAVFDKNLYDKLV</u> <u>SSFLEMKSEK</u>

NHIBIT tag in bold, Linker in italics, Mac1 underlined

A PCR product encoding SARS Cov-2 macrodomain residues 1 to 176 was cloned into the HindIII and XhoI restriction sites of the pBIT3.1-C vector(Promega) which has an C-terminal 11 amino acid HiBiT tag.

PBIT CHIBIT SARS-CoV-2 Macrodomain

MG<u>PEVNSFSGYLKLTDNVYIKNADIVEEAKKVKPTVVVNAANVYLKHGGGVAGALNKATNN</u> <u>AMQVESDDYIATNGPLKVGGSCVLSGHNLAKHCLHVVGPNVNKGEDIQLLKSAYENFNQH</u> <u>EVLLAPLLSAGIFGADPIHSLRVCVDTVRTNVYLAVFDKNLYDKLVSSFLEMKSEK</u>*GSSGGS SG***VSGWRLFKKIS**

CHIBIT tag in bold, Linker in italics, Mac 1 underlined

Both constructs were evaluated for CETSA experiments and found to give similar results. For the results described the CHIBIT tag was used.

HiBiT Cellular Thermal Shift Assay (HiBiT-CETSA). To assess the engagement of nsp3-mac1 in intact cells, a HiBiT Cellular Thermal Shift Assay was conducted as described by Ramachandran et al. (*23*). The Nano-Glo HiBiT lytic detection system (Promega) was used according to the manufacturer’s instructions. Briefly, HEK293 cells were reverse-transfected with HiBiT-tagged nsp3-mac1 at a concentration of 400,000 cells/mL using the FuGENE transfection reagent (Promega). Following a 24-hour incubation period, the cells were trypsinized and resuspended in assay medium at the same concentration.

For CETSA, the cells were divided into two groups: one treated with 30 μM of the compounds and the other with DMSO as a control. For ITDRF CETSA, cells were treated with varying concentrations of the compounds as indicated. Ten microliters of the cell suspension were seeded into each well of a 384-well white PCR plate and incubated for 1 hour at 37°C with 5% CO2 in a humidified incubator. After incubation, the samples were subjected to heating in a PCR thermocycler at the indicated temperature gradient (CETSA) or a fixed temperature of 50.6°C (ITDRF CETSA). The plate was then left to stand at room temperature for 5 minutes, after which 10 μL of Nano-Glo HiBiT lytic mix was added to each well. The luminescence signals were measured using a PHERAstar FSX and analyzed with GraphPad Prism software (v.9 or v.10).

### Protein expression and purification for biochemical assay

The expression and purification for the biochemical assay was described here in detail dx.doi.org/10.17504/protocols.io.4r3l2qb9jl1y/v1.

### Structural Biology

#### Protein expression and purification for crystallisation

Constructs for initial crystallography (P4_3_ form) and assay were from Dr. M Schuller as detailed in ref (*13*) and described in the following protocol dx.doi.org/10.17504/protocols.io.dm6gpz9xplzp/v1. The second crystallography construct (P1 2_1_ 1 form) was cloned into a pNIC vector with a His Sumo tag (Addgene: 215810) and residues (2 – 170) for bacterial expression described in detail in the following protocol dx.doi.org/10.17504/protocols.io.n92ld8jm8v5b/v1. All constructs were transformed into BL21(DE3)R3-pRARE2-RR for expression.

Cultures in LB media were used to inoculate either TB media and induced with 0.4 mM isopropyl 1-thio-β-D-galactopyranoside overnight at 18°C (P4_3_ form) or autoinduction TB media and left to induce via autoinduction, reducing the temperature when the cells reached OD2 to 18°C overnight (P1 2_1_ 1 form and assay construct).

Cells were lysed by sonication and all proteins were purified by nickel affinity chromatography. The crystallography proteins were then desalted and cleaved overnight. The assay construct was not cleaved. The tag on the P4_3_ construct was cleaved using tobacco etch protease (Tev) while the tag on the P1 2_1_ 1 construct was cleaved using SENP1. Both crystallography proteins were then purified by reverse nickel affinity chromatography to remove the cleaved tag. All proteins were concentrated and purified by size exclusion chromatography using a HiLoad 16/60 Superdex 75 pg gel filtration column. The mass of the eluted samples were confirmed by mass spec before being concentrated to 21.6 mg/ml and flash frozen in 10 mM HEPES pH7.5, 0.5M NaCl, 5% glycerol, 0.5 mM TCEP (p1 21 1 construct) or 50 mM HEPES pH7.5, 0.5M NaCl, 5% glycerol, 0.5 mM TCEP (p43 construct). The assay construct was concentrated to 6.2 mg/ml and flash frozen in 50 mM HEPES pH7.5, 0.5M NaCl, 5% glycerol, 0.5 mM TCEP.

#### Crystallisation of SARS-CoV-2 nsp3-mac1 in P4_3_

Crystallisation of nsp3-mac1 was performed as previously described (13). In shorts, crystals were grown using the sitting drop vapour diffusion method in SwissCI 3 lens crystallisation plates using a SPT mosquito liquid handler. The crystallisation solution consisted of 100 mM CHES (pH 9.5) and 30% (w/v) PEG3000. Each drop contained 150 nl of protein solution (47 mg/ml) and 150 nl reservoir solution. Plates were incubated at 20°C and monitored using a Formulatrix Rock Imager. (dx.doi.org/10.17504/protocols.io.e6nvw1qb2lmk/v1)

#### Crystallisation of SARS-CoV-2 nsp3 macrodomain in P1 2_1_ 1

To obtain crystals in P1 21 1, nsp3-mac1 was used at 21.6 mg/ml. Crystallisation was performed same as described above but the crystallisation solution consisted of 0.1 M MES (pH 6.5) and 30% (w/v) PEG3000 and each drop contained 400 nl protein solution, 300 nl reservoir solution, and 100 nl seed stock (1:2 dilution), for a final volume of 800 nl (dx.doi.org/10.17504/protocols.io.261ge5ej7g47/v1).

#### Sequence of SARS-CoV-2 nsp3 macrodomain (P1 2_1_ 1 form)

Vector: pNIC-NOTAG-GG (Addgene: 215810)

With tag:

MGSSHHHHHHMASMSDSEVNQEAKPEVKPEVKPETHINLKVSDGSSEIFFKIKKTTPLRRLMEAFAKRQGKEMDSLRFLYDGIRIQADQTPEDLDMEDNDIIEAHREQIGGGEVNSFSGYLKLTDNVYIKNADIVEEAKKVKPTVVVNAANVYLKHGGGVAGALNKATNNAMQVESDDYIATNGPLKVGGSCVLSGHNLAKHCLHVVGPNVNKGEDIQLLKSAYENFNQHEVLLAPLLSAGIFGADPIHSLRVCVDTVRTNVYLAVFDKNLYDKLVSSFLE

Without tag:

GEVNSFSGYLKLTDNVYIKNADIVEEAKKVKPTVVVNAANVYLKHGGGVAGALNKATNNAMQVESDDYIATNGPLKVGGSCVLSGHNLAKHCLHVVGPNVNKGEDIQLLKSAYENFNQHEVLLAPLLSAGIFGADPIHSLRVCVDTVRTNVYLAVFDKNLYDKLVSSFLE

#### Compound soaking

Compound soaking was done as described previously (23). Compounds were added to the drops containing crystals using an Echo liquid handling system (Beckman Coulter) (final concentration, 10% (v/v) DMSO) and were incubated for 1 h before the crystals were harvested and cryo-cooled in liquid nitrogen.

#### Co-crystallisation of SARS-CoV-2 nsp3 macrodomain with ligands in P1 2_1_ 1

For co-crystallisation, 40 nl of compound (∼5% of final drop volume) was first dispensed to the lenses of a SwissCI 3 lens crystallisation plate using the Echo liquid handling system, before protein, reservoir and seed solution was added as described previously (dx.doi.org/10.17504/protocols.io.n2bvjnb9bgk5/v1).

#### Data collection, processing and model building

Data was collected at the I04-1 beamline (Diamond Light Source, UK) at 100 K and automatically processed with Diamond Light Source’s autoprocessing pipelines using XDS (*24*) and either xia (*25*), autoPROC (*26*) or DIALS (*27*) with the default settings. DIMPLE (*28*) was used to generate the electron density maps. Ligand restraints were calculated with GRADE (*29*) and ligand-binding events were identified using PanDDA (*30*) or PanDDA2 (*31*) if electron density for ligand was not already clearly visible in the DIMPLE map. Iterative model building was done manually in Coot (*32*) and structures were refined using BUSTER (*33*). Coordinates and structure factors for the ligand-bound SARS-CoV-2 nsp3-mac1 structures are deposited in the Protein Data Bank (group deposition G_1002283).

## Supporting information

Bioactivity data

## Supplementary Materials

We disclose bioactivity data on over 1000 compounds tested as part of the nsp3-mac1 inhibitor discovery efforts.

## Acknowledgment

This work was supported by the NIH and NIAID Antiviral Drug Discovery (AViDD) grant (U19AI171399). The content is solely the responsibility of the authors and does not necessarily represent the official views of the National Institute of Health.

## References

1. A. von Delft, M. D. Hall, A. D. Kwong, L. A. Purcell, K. S. Saikatendu, U. Schmitz, J. A. Tallarico, A. A. Lee, Accelerating antiviral drug discovery: lessons from COVID-19. Nat. Rev. Drug Discov. 22, 585–603 (2023).

2. B. Tan, X. Zhang, A. Ansari, P. Jadhav, H. Tan, K. Li, A. Chopra, A. Ford, X. Chi, F. X. Ruiz, E. Arnold, X. Deng, J. Wang, Design of a SARS-CoV-2 papain-like protease inhibitor with antiviral efficacy in a mouse model. Science 383, 1434–1440 (2024).

3. M. R. Garnsey, M. C. Robinson, L. T. Nguyen, R. Cardin, J. Tillotson, E. Mashalidis, A. Yu, L. Aschenbrenner, A. Balesano, A. Behzadi, B. Boras, J. S. Chang, H. Eng, A. Ephron, T. Foley, K. K. Ford, J. M. Frick, S. Gibson, L. Hao, B. Hurst, A. S. Kalgutkar, M. Korczynska, Z. Lengyel-Zhand, L. Gao, H. R. Meredith, N. C. Patel, J. Polivkova, D. Rai, C. R. Rose, H. Rothan, S. K. Sakata, T. R. Vargo, W. Qi, H. Wu, Y. Liu, I. Yurgelonis, J. Zhang, Y. Zhu, L. Zhang, A. A. Lee, Discovery of SARS-CoV-2 papain-like protease (PLpro) inhibitors with efficacy in a murine infection model, bioRxiv (2024)p. 2024.01.26.577395.

4. L. C. Russo, R. Tomasin, I. A. Matos, A. C. Manucci, S. T. Sowa, K. Dale, K. W. Caldecott, L. Lehtiö, D. Schechtman, F. C. Meotti, A. Bruni-Cardoso, N. C. Hoch, The SARS-CoV-2 Nsp3 macrodomain reverses PARP9/DTX3L-dependent ADP-ribosylation induced by interferon signaling. J. Biol. Chem. 297, 101041 (2021).

5. J. G. M. Rack, D. Perina, I. Ahel, Macrodomains: Structure, Function, Evolution, and Catalytic Activities. Annu. Rev. Biochem. 85, 431–454 (2016).

6. A. R. Fehr, G. Jankevicius, I. Ahel, S. Perlman, Viral Macrodomains: Unique Mediators of Viral Replication and Pathogenesis. Trends Microbiol. 26, 598–610 (2018).

7. J. G. M. Rack, V. Zorzini, Z. Zhu, M. Schuller, D. Ahel, I. Ahel, Viral macrodomains: a structural and evolutionary assessment of the pharmacological potential. Open Biol. 10, 200237 (2020).

8. Y. M. Alhammad, S. Parthasarathy, R. Ghimire, C. M. Kerr, J. J. O’Connor, J. J. Pfannenstiel, D. Chanda, C. A. Miller, N. Baumlin, M. Salathe, R. L. Unckless, S. Zuñiga, L. Enjuanes, S. More, R. Channappanavar, A. R. Fehr, SARS-CoV-2 Mac1 is required for IFN antagonism and efficient virus replication in cell culture and in mice. Proc. Natl. Acad. Sci. U. S. A. 120, e2302083120 (2023).

9. T. Y. Taha, R. K. Suryawanshi, I. P. Chen, G. J. Correy, M. McCavitt-Malvido, P. C. O’Leary, M. P. Jogalekar, M. E. Diolaiti, G. R. Kimmerly, C.-L. Tsou, R. Gascon, M. Montano, L. Martinez-Sobrido, N. J. Krogan, A. Ashworth, J. S. Fraser, M. Ott, A single inactivating amino acid change in the SARS-CoV-2 NSP3 Mac1 domain attenuates viral replication in vivo. PLoS Pathog. 19, e1011614 (2023).

10. L. S. Voth, J. J. O’Connor, C. M. Kerr, E. Doerger, N. Schwarting, P. Sperstad, D. K. Johnson, A. R. Fehr, Unique Mutations in the Murine Hepatitis Virus Macrodomain Differentially Attenuate Virus Replication, Indicating Multiple Roles for the Macrodomain in Coronavirus Replication. J. Virol. 95, e0076621 (2021).

11. R. Abraham, D. Hauer, R. L. McPherson, A. Utt, I. T. Kirby, M. S. Cohen, A. Merits, A. K. L. Leung, D. E. Griffin, ADP-ribosyl-binding and hydrolase activities of the alphavirus nsP3 macrodomain are critical for initiation of virus replication. Proc. Natl. Acad. Sci. U. S. A. 115, E10457–E10466 (2018).

12. R. Abraham, R. L. McPherson, M. Dasovich, M. Badiee, A. K. L. Leung, D. E. Griffin, Both ADP-Ribosyl-Binding and Hydrolase Activities of the Alphavirus nsP3 Macrodomain Affect Neurovirulence in Mice. MBio 11 (2020).

13. M. Schuller, G. J. Correy, S. Gahbauer, D. Fearon, T. Wu, R. E. Díaz, I. D. Young, L. Carvalho Martins, D. H. Smith, U. Schulze-Gahmen, T. W. Owens, I. Deshpande, G. E. Merz, A. C. Thwin, J. T. Biel, J. K. Peters, M. Moritz, N. Herrera, H. T. Kratochvil, QCRG Structural Biology Consortium, A. Aimon, J. M. Bennett, J. Brandao Neto, A. E. Cohen Dias, A. Douangamath, L. Dunnett, O. Fedorov, M. P. Ferla, M. R. Fuchs, T. J. Gorrie-Stone, J. M. Holton, M. G. Johnson, T. Krojer, G. Meigs, A. J. Powell, J. G. M. Rack, V. L. Rangel, S. Russi, R. E. Skyner, C. A. Smith, A. S. Soares, J. L. Wierman, K. Zhu, P. O’Brien, N. Jura, A. Ashworth, J. J. Irwin, M. C. Thompson, J. E. Gestwicki, F. von Delft, B. K. Shoichet, J. S. Fraser, I. Ahel, Fragment binding to the Nsp3 macrodomain of SARS-CoV-2 identified through crystallographic screening and computational docking. Sci Adv 7 (2021).

14. S. Gahbauer, G. J. Correy, M. Schuller, M. P. Ferla, Y. U. Doruk, M. Rachman, T. Wu, M. Diolaiti, S. Wang, R. J. Neitz, D. Fearon, D. S. Radchenko, Y. S. Moroz, J. J. Irwin, A. R. Renslo, J. C. Taylor, J. E. Gestwicki, F. von Delft, A. Ashworth, I. Ahel, B. K. Shoichet, J. S. Fraser, Iterative computational design and crystallographic screening identifies potent inhibitors targeting the Nsp3 macrodomain of SARS-CoV-2. Proc. Natl. Acad. Sci. U. S. A. 120, e2212931120 (2023).

15. S. Wazir, T. A. O. Parviainen, J. J. Pfannenstiel, M. T. H. Duong, D. Cluff, S. T. Sowa, A. Galera-Prat, D. Ferraris, M. M. Maksimainen, A. R. Fehr, J. P. Heiskanen, L. Lehtiö, Discovery of 2-Amide-3-methylester Thiophenes that Target SARS-CoV-2 Mac1 and Repress Coronavirus Replication, Validating Mac1 as an Antiviral Target. J. Med. Chem. 67, 6519–6536 (2024).

16. G. Chessari, R. Grainger, R. S. Holvey, R. F. Ludlow, P. N. Mortenson, D. C. Rees, C-H functionalisation tolerant to polar groups could transform fragment-based drug discovery (FBDD). Chem. Sci. 12, 11976–11985 (2021).

17. H. S. Yu, K. Modugula, O. Ichihara, K. Kramschuster, S. Keng, R. Abel, L. Wang, General Theory of Fragment Linking in Molecular Design: Why Fragment Linking Rarely Succeeds and How to Improve Outcomes. J. Chem. Theory Comput. 17, 450–462 (2021).

18. Y. Unoh, S. Uehara, K. Nakahara, H. Nobori, Y. Yamatsu, S. Yamamoto, Y. Maruyama, Y. Taoda, K. Kasamatsu, T. Suto, K. Kouki, A. Nakahashi, S. Kawashima, T. Sanaki, S. Toba, K. Uemura, T. Mizutare, S. Ando, M. Sasaki, Y. Orba, H. Sawa, A. Sato, T. Sato, T. Kato, Y. Tachibana, Discovery of S-217622, a Noncovalent Oral SARS-CoV-2 3CL Protease Inhibitor Clinical Candidate for Treating COVID-19. J. Med. Chem. 65, 6499– 6512 (2022).

19. E. Kojima, A. Iimuro, M. Nakajima, H. Kinuta, N. Asada, Y. Sako, Z. Nakata, K. Uemura, S. Arita, S. Miki, C. Wakasa-Morimoto, Y. Tachibana, Pocket-to-Lead: Structure-Based De Novo Design of Novel Non-peptidic HIV-1 Protease Inhibitors Using the Ligand Binding Pocket as a Template. J. Med. Chem. 65, 6157–6170 (2022).

20. S. Yoshida, Y. Sako, E. Nikaido, T. Ueda, I. Kozono, Y. Ichihashi, A. Nakahashi, M. Onishi, Y. Yamatsu, T. Kato, J. Nishikawa, Y. Tachibana, Peptide-to-Small Molecule: Discovery of Non-Covalent, Active-Site Inhibitors of β-Herpesvirus Proteases. ACS Med. Chem. Lett. 14, 1558–1566 (2023).

21. W. McCorkindale, I. Ahel, H. Barr, G. J. Correy, J. S. Fraser, N. London, M. Schuller, K. Shurrush, A. A. Lee, Fragment-Based Hit Discovery via Unsupervised Learning of Fragment-Protein Complexes, bioRxiv (2022)p. 2022.11.21.517375.

22. A. Gaulton, L. J. Bellis, A. P. Bento, J. Chambers, M. Davies, A. Hersey, Y. Light, S. McGlinchey, D. Michalovich, B. Al-Lazikani, J. P. Overington, ChEMBL: a large-scale bioactivity database for drug discovery. Nucleic Acids Res. 40, D1100–D1107 (2011).

23. S. Ramachandran, M. Szewczyk, S. H. Barghout, A. Ciulli, D. Barsyte-Lovejoy, V. Vu, HiBiT Cellular Thermal Shift Assay (HiBiT CETSA). Methods Mol. Biol. 2706, 149–165 (2023).

24. W. Kabsch, XDS. Acta Crystallogr. D Biol. Crystallogr. 66, 125–132 (2010).

25. G. Winter, C. M. C. Lobley, S. M. Prince, Decision making in xia2. Acta Crystallogr. D Biol. Crystallogr. 69, 1260–1273 (2013).

26. C. Vonrhein, C. Flensburg, P. Keller, A. Sharff, O. Smart, W. Paciorek, T. Womack, G. Bricogne, Data processing and analysis with the autoPROC toolbox. Acta Crystallogr. D Biol. Crystallogr. 67, 293–302 (2011).

27. G. Winter, D. G. Waterman, J. M. Parkhurst, A. S. Brewster, R. J. Gildea, M. Gerstel, L. Fuentes-Montero, M. Vollmar, T. Michels-Clark, I. D. Young, N. K. Sauter, G. Evans, DIALS: implementation and evaluation of a new integration package. Acta Crystallogr D Struct Biol 74, 85–97 (2018).

28. M. Wojdyr, R. Keegan, G. Winter, DIMPLE-a pipeline for the rapid generation of difference maps from protein crystals with putatively bound ligands. Acta Crystallogr. (2013).

29. O. S. Smart, J. H. A. Sharff, T. O. Womack, C. Flensburg, P. Keller, W. Paciorek, C. Vonrhein, G. Bricogne, Grade2 Version 1.6.0. Cambridge, United Kingdom: Global Phasing Ltd (2021).

30. N. M. Pearce, T. Krojer, A. R. Bradley, P. Collins, R. P. Nowak, R. Talon, B. D. Marsden, S. Kelm, J. Shi, C. M. Deane, F. von Delft, A multi-crystal method for extracting obscured crystallographic states from conventionally uninterpretable electron density. Nat. Commun. 8, 15123 (2017).

31. C. ConorFWild, ConorFWild/pandda_2_gemmi: Alpha (https://zenodo.org/records/11035775).

32. P. Emsley, B. Lohkamp, W. G. Scott, K. Cowtan, Features and development of Coot. Acta Crystallogr. D Biol. Crystallogr. 66, 486–501 (2010).

33. G. Bricogne, E. Blanc, M. Brandl, C. Flensburg, P. Keller, W. Paciorek, P. Roversi, A. Sharff, O. S. Smart, C. Vonrhein, T. O. Womack, BUSTER. Cambridge, United Kingdom: Global Phasing Ltd (2017).

